# HuR-dependent expression of RyR2 contributes to calcium-mediated thermogenesis in brown adipocytes

**DOI:** 10.1101/2024.10.22.619637

**Authors:** Adrienne R. Guarnieri, Sarah R. Anthony, Pooja Acharya, Bo Yao Wen, Lindsey Lanzillotta, Rylie Gavin, Michael Tranter

## Abstract

Several uncoupling protein 1 (UCP1)-independent thermogenic pathways have been described in thermogenic adipose tissue, including calcium-mediated thermogenesis in beige adipocytes via sarco/endoplasmic reticulum ATPase (SERCA). We have previously shown that adipocyte-specific deletion of the RNA binding protein human antigen R (HuR) results in thermogenic dysfunction independent of UCP1 expression. RNA sequencing revealed the downregulation of several genes involved in calcium ion transport upon HuR deletion. The goal of this work was to define the HuR-dependent mechanisms of calcium driven thermogenesis in brown adipocytes.

We generated (BAT)-specific HuR-deletion (BAT-HuR^-/-^) mice and show that their body weight, glucose tolerance, brown and white adipose tissue weights, and total lipid droplet size were not significantly different compared to wild-type. Similar to our initial findings in Adipo-HuR^-/-^ mice, mice with BAT-specific HuR deletion are cold intolerant following acute thermal challenge at 4°C, demonstrating specificity of acute HuR-dependent thermogenesis to BAT. We also found decreased expression of ryanodine receptor 2 (RyR2), but no changes in RyR2, SERCA1, SERCA2, or UCP1 expression, in BAT from BAT-HuR^-/-^ mice. Next, we used Fluo-4 calcium indicator dye to show that genetic deletion or pharmacological inhibition of HuR blunts the increase in cytosolic calcium concentration in SVF-derived primary brown adipocytes. Moreover, we saw a similar blunting in β-adrenergic-mediated heat generation, as assessed by ERtherm AC fluorescence, in SVF-derived brown adipocytes following HuR inhibition or deletion. Mechanistically, we show that HuR directly binds and reduces the decay rate of RyR2 mRNA in brown adipocytes, and stabilization of RyR2 via S107 rescues β-adrenergic-mediated cytosolic calcium increase and heat generation in HuR deficient brown adipocytes.

In conclusion, our results suggest that HuR-dependent control of RyR2 expression plays a significant role in the thermogenic function of brown adipose tissue through modulation of SR calcium cycling.

## Introduction

Obesity is defined as an excess accumulation of body fat and is most commonly driven by imbalances between energy consumption and energy expenditure that, resulting in the storage of excess energy within adipose tissue (AT) depots throughout the body. Obesity represents a growing issue worldwide where its prevalence has progressively increased over the past few decades.^1,2^ Currently over 100 million children (∼5% of global population) and 600 million adults (∼12% global population) are affected by obesity.^3,4^ This rise is particularly evident in America where the age-adjusted prevalence has grown from 30.5% at the turn of the century, to 42.4% in 2018.^4,5^

The therapeutic potential of increasing whole-body energy expenditure to mitigate obesity development was highlighted by the identification of inducible, metabolically active brown adipose tissue in adult humans.^6–8^ In contrast to white adipose tissue which functions for lipid storage, mitochondria-heavy brown and beige adipose tissue function primarily for energy dissipation in the form of heat through non-shivering thermogenesis (NST). NST is activated by the sympathetic stimulation of β-adrenergic receptors on the adipocyte cell surface and is primarily driven by action of the mitochondrial inner membrane protein uncoupling protein 1 (UCP1).^9^ However, the existence of UCP1-independent thermogenic pathways was suggested after studies reported UCP1-deficient mice housed at room temperature were surprisingly protected from diet-induced obesity.^10–12^

Calcium cycling was observed in brown adipocytes as early as the 1970s, but the involvement of calcium transport in thermogenesis wasn’t implicated until much later when it was shown that elevation of cytosolic calcium, originating from an intracellular source, was sufficient to potentiate adrenergic-mediated thermogenesis.^13,14^ Calcium-dependent thermogenesis is mediated by action of sarcoendoplasmic-reticulum (SR) calcium ATPase (SERCA), which hydrolyzes ATP to transport cytosolic calcium into the SR, and Ryanodine Receptor (RyR), which releases sequestered SR calcium to the cytosol.^15,16^ More recently, Ikeda et. al demonstrated SR calcium cycling as a potent mediator of thermogenic metabolism in beige adipocytes.^17^

Our lab has recently identified the RNA binding protein Human Antigen R (HuR), as a mediator of adipose tissue thermogenesis.^18^ Specifically, our results showed a reduced ability of mice with an adipose tissue specific deletion of HuR (Adipo-HuR^-/-^) to thermoregulate in response to acute cold challenge at 4°C, concomitant with a HuR-dependent decrease in the expression of calcium handling genes in BAT.^18^ Thus, the goal of this work was to utilize BAT-specific HuR deletion mice (BAT-HuR^-/-^) to determine the mechanistic and functional impact of HuR on BAT-mediated SR calcium cycling and thermogenesis.

## Methods

### Animals and *In Vivo* Experiments

All mouse studies were approved by the University of Cincinnati and The Ohio State University Institutional Animal Care and Use Committee (IACUC). HuR-floxed mice described by Ghosh et. al^19^ were obtained from Jackson labs (stock #: 021431) and crossed with mice harboring a UCP1-specific Cre recombinase transgene (UCP1-Cre; Jackson labs, stock #: 024670) to generate a BAT-specific deletion of HuR (BAT-HuR^-/-^). Wild-type *(wt/wt-cre*^*+*^; *flox/flox-cre*^*-*^; *wt/flox-cre*^*-*^) littermates were used as controls. Wild-type C57BL/J mice (Jackson labs, stock #: 010803) were used to establish cell cultures for experiments involving pharmacological inhibition of HuR. All studies include equal proportions of male and female mice housed at 23°C.

For high fat diet studies, 9-week-old mice were placed on a high fat diet (HFD; 45% kcal from fat; Research Diets, Inc) or control chow (10% kcal from fat) for 12 weeks, and total body weight was assessed weekly. Glucose tolerance was assessed on 10-12-week-old mice on control chow diet following an intraperitoneal injection of 45% glucose solution (1 g/kg).

For the acute cold challenge, mice were individually housed with food and bedding removed at 4°C and core body temperature was monitored via rectal probe thermometer. Mice were returned to room temperature when core body temperature fell outside of euthermic range (below 32°C), or after 6 hours of cold exposure.

### Cell Culture and Pharmacology

For primary adipocyte cell culture, mice 6-12 weeks of age were sedated with isoflurane and euthanized via bilateral pneumothorax followed immediately by adipose tissue extraction. Brown adipocyte cells were established from the stromal vascular fraction of isolated brown adipose tissue by enzymatic digestion with collagen I, and differentiation was induced by growth in medium containing 2% FBS, 40% MCDB201, 1x L-Ascorbic Acid, 1x ITS MIX, 0.1 mM L-ascorbic acid-2-phosphate, and 1 nM of dexamethasone. Three days after induction, the culture medium was additionally supplemented with 5 μg/mL insulin, 1 nM T3, 500 nM isobutylmethylxanthine (IBMX), 1 μM dexamethasone, and 50 μM indomethacin. Maturation of brown adipocytes was confirmed via visualization of multilocular lipid deposits upon oil red O staining and qPCR assessment of brown adipose specific genes.

For HuR inhibition studies, differentiated brown adipocytes from control mice were treated overnight (12-16 hours) with 2.5 μM Dihydrotanshinone-I (DHTS). To stimulate SR calcium release and thermogenesis, brown adipocytes were treated with 10 μM Isoproterenol (ISO) or 10 μM CL316,243 (CL) to stimulate β-AR and β3-AR-specific signaling, respectively. To assess the effects of RyR stabilization, cells were treated with 10 μM of S107 for 10 minutes prior to imaging.

### Calcium and heat imaging

Differentiated mature brown adipocytes were imaged for cytosolic calcium using Fluo-4 Calcium Assay kit (Invitrogen F10471). Cells were incubated with 4 μM Fluo-4 AM dye in HHBS buffer for 45 minutes at 37°C. After removal of excess extracellular Fluo-4 with a PBS wash, cells were imaged on a Stellaris 8 confocal microscope and fluorescence intensity was assessed at excitation/emission wavelengths of 490/515 using a BioTek Cytation 5. Changes in cytosolic calcium were determined by comparing ISO-stimulated values to basal Fluo-4 fluorescence (F/F_o_).

Cellular heat production was determined using ERtherm AC dye as previously described.^20^ Differentiated brown adipocytes were incubated with 250 nm ERtherm AC in Phenol red-free DMEM for 30 minutes at 25°C to allow for equilibration. Basal fluorescent values were measured at excitation/emission wavelengths of 555/655 nm using BioTek Cytation 5. Cells were then stimulated with 10 μM CL316,243 and fluorescence intensity was recorded every hour. Heat production quenches fluorescence. Quantification of cellular heat production is represented as the reciprocal of CL-stimulated values over basal fluorescent values [1/(F/F_o_)].

### RNA isolation and mRNA assessment

RNA was isolated from BAT and cDNA was synthesized for qPCR using iScript RT Supermix (Bio-Rad). Samples were run on a Bio-Rad CFX96 Touch to assess mRNA levels of gene transcripts (see table 2.1 for primer sequences). Nuclear and mitochondrial DNA were isolated using a NucleoSpin Tissue gDNA isolation kit (Takara Bio 740952) per manufacturer’s instructions and nuclear and mitochondrial DNA was quantified via qPCR assessment of 28S and COX1 levels, respectively.

**Table 1.**
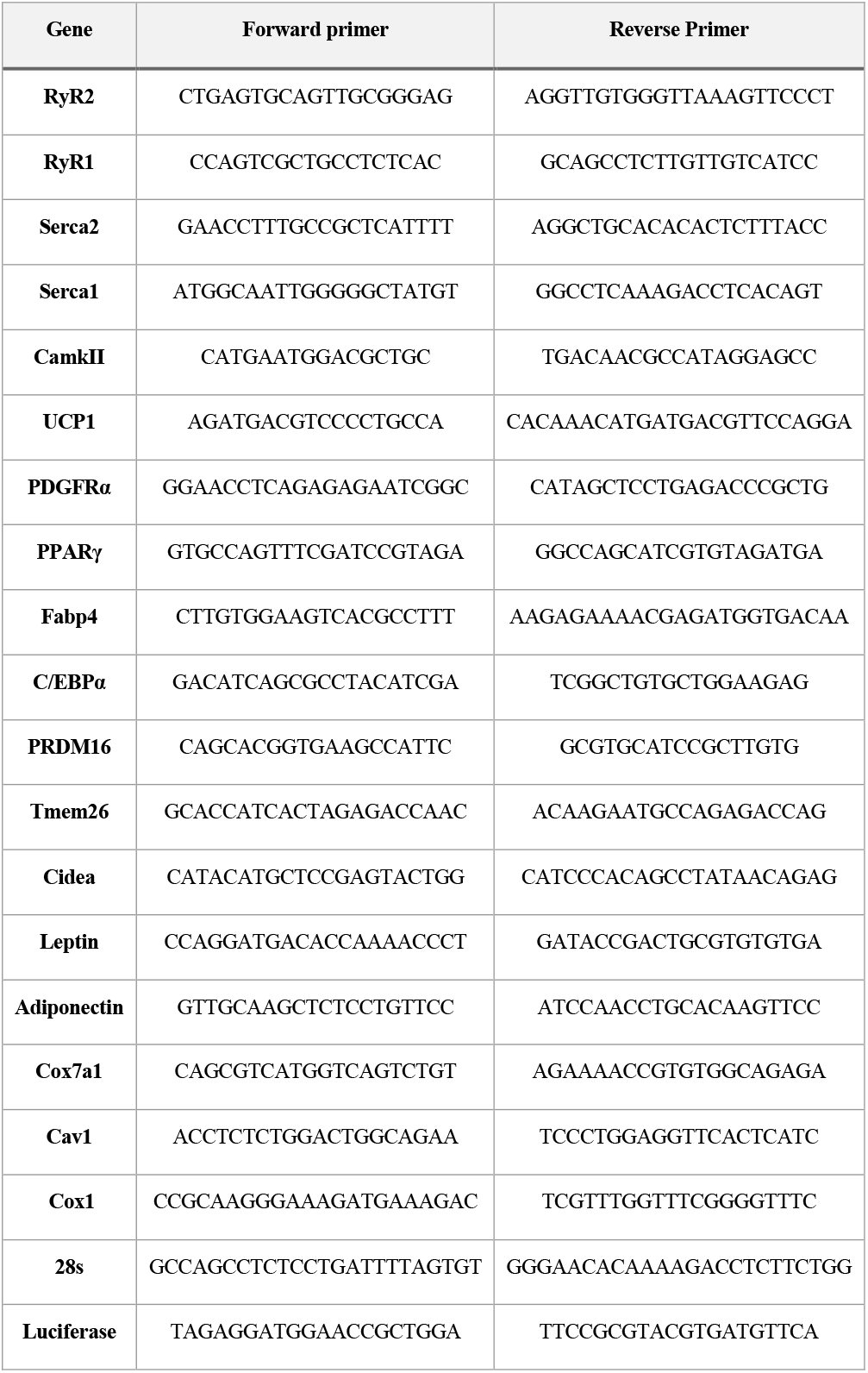
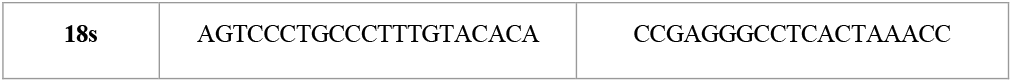
Primer sequences for qPCR.

For RNA cross-linking and immunoprecipitation (CLIP) experiments, cells were grown in 10cm culture dishes and then UV cross-linked at 200,000 μJ/cm^3^ at 240nm wavelength. Cells were then lysed via dounce homogenization and lysate was added to 0.75 mg prewashed Protein G Dynabeads (Invitrogen 10004D) in conjunction with 4 ug of HuR antibody (AbCam ab200342) as previously described by our lab.^21^ RNA was eluted and lysate samples were spiked with 5 ng of Luciferase control RNA (Promega L4561) as a housekeeping control for total RNA isolation efficiency in qPCR assessment of HuR-bound target genes.

To assess mRNA transcript stability, SVF-derived brown adipocytes were treated with 2.5 μg/ml Actinomycin D and 2.5 μM vehicle or DHTS as we have previously described.^22^

### Protein isolation and quantification

Protein extraction from adipose tissue was done via homogenization in RIPA buffer with 0.5 mM DTT, 0.2 mM sodium-orthovanadate, and a protease inhibitor (1:1000; Invitrogen 78430) followed by centrifugation at 16,000 RPM for 10 minutes for separation of protein fraction from the mature adipocyte layer. Nuclear and cytoplasmic fractions were collected by lysing cells directly in ice cold Solution A (10mM HEPES pH 9, 1.5 mM MgCl_2_, 10 mM KCl, 0.5 mM DTT, 0.1% Triton X, and protease inhibitor cocktail) followed by dounce homogenization and incubation on ice for 10 minutes. The lysate was then centrifuged at 5000xG for 10 minutes, and the supernatant was saved as cytoplasmic fraction, while the pellet was resuspended in Solution C (20 mM HEPES pH 7.9, 25% glycerol, 0.6 M KCl, 1.5 mM MgCl_2_, 0.2 mM EDTA, 0.5 mM DTT, and protease inhibitor cocktail), incubated on ice for 40 minutes with vortexing every 10 minutes, and centrifuged at 10000xG for 15 minutes. The supernatant was then retained as the nuclear fraction.

Protein extract (15 μg total; 10 μg nuclear; 30 μg cytoplasmic) was separated on a 10% polyacrylamide gel and transferred to a nitrocellulose membrane. Blocking was performed for 1 h at room temperature using 4% BSA in 0.1% Tween 20, tris-buffered saline (T-TBS). Primary antibodies for HuR (AbCam ab200342), UCP1 (Novus Biologicals NBP2-20,796), and RyR2 (Invitrogen PA5-87416) were incubated overnight at 4°C, and appropriate secondary antibodies were incubated for 1 h at room temperature in T-TBS. Loading was normalized to total protein using TGX Stain Free precast gels (BioRad, Hercules, CA) as described.^444^ Images were captured and analyzed using a ChemiDoc Imaging System and ImageLab software.

### Histology

Tissues were fixed using 4% paraformaldehyde, paraffin embedded by the Cincinnati Children’s Hospital Medical Center Department of Pathology Research Core (Cincinnati, OH, USA), and sectioned at 5 μm thickness prior to staining with hematoxylin and eosin (H&E). Lipid droplet size was calculated using edge thresholding in ImageJ (NIH, Bethesda, MD). Immunofluorescent staining for RyR2 was performed using an Alexa Fluor conjugated antibody kit (Invitrogen Z25351) with 10 ug of conjugated primary antibody for 1 hour at 37°C, washed with TBS-T, and mounted with DAPI for visualization of nuclei. Images were obtained on a Stellaris 8 confocal microscope and mean fluorescence intensity was quantified for each individual cell.

### Statistical analysis

Data is represented as mean ± SEM unless otherwise noted. All data was tested for normality (Shapiro-Wilk) and equal variance (Brown-Forsythe). Individual comparisons were done using Student’s t-test. Comparisons between multiple groups were performed using a one-way ANOVA. Serial measurements were analyzed using multiple comparisons within a two-way ANOVA. Cold challenge results were assessed using a log-rank (Mantel-Cox) test. All graphing and statistical analyses were performed using GraphPad Prism 10.

## Results

### Generation and phenotypic characterization of BAT-specific HuR deletion mice

Our previous work examining HuR deletion across all adipose tissue using an adiponectin promoter Cre recombinase (Adipo-HuR^-/-^) showed that HuR is a mediator of adipose tissue thermogenesis. To confirm the specificity of this observation to HuR activity in BAT and rule out compensatory impacts on thermogenic capacity driven by the white adipose tissue, we generated brown adipose-specific HuR deletion mice (BAT-HuR^-/-^) using a UCP1-Cre promoter. As expected, we observed ∼80% reduction in HuR protein in BAT (Fig, 1A-B), but no change in HuR protein levels in white adipose tissue (Fig. 1A, C-D). Additionally, BAT-HuR^-/-^ mice exhibited no differences in the BAT mRNA expression of brown adipogenesis driver, PRDM16 (Fig. 1E), nor were there any differences in brown adipose tissue lipid droplet size or glucose tolerance from wildtype mice (Fig. 1F-H).

**Figure 1.**
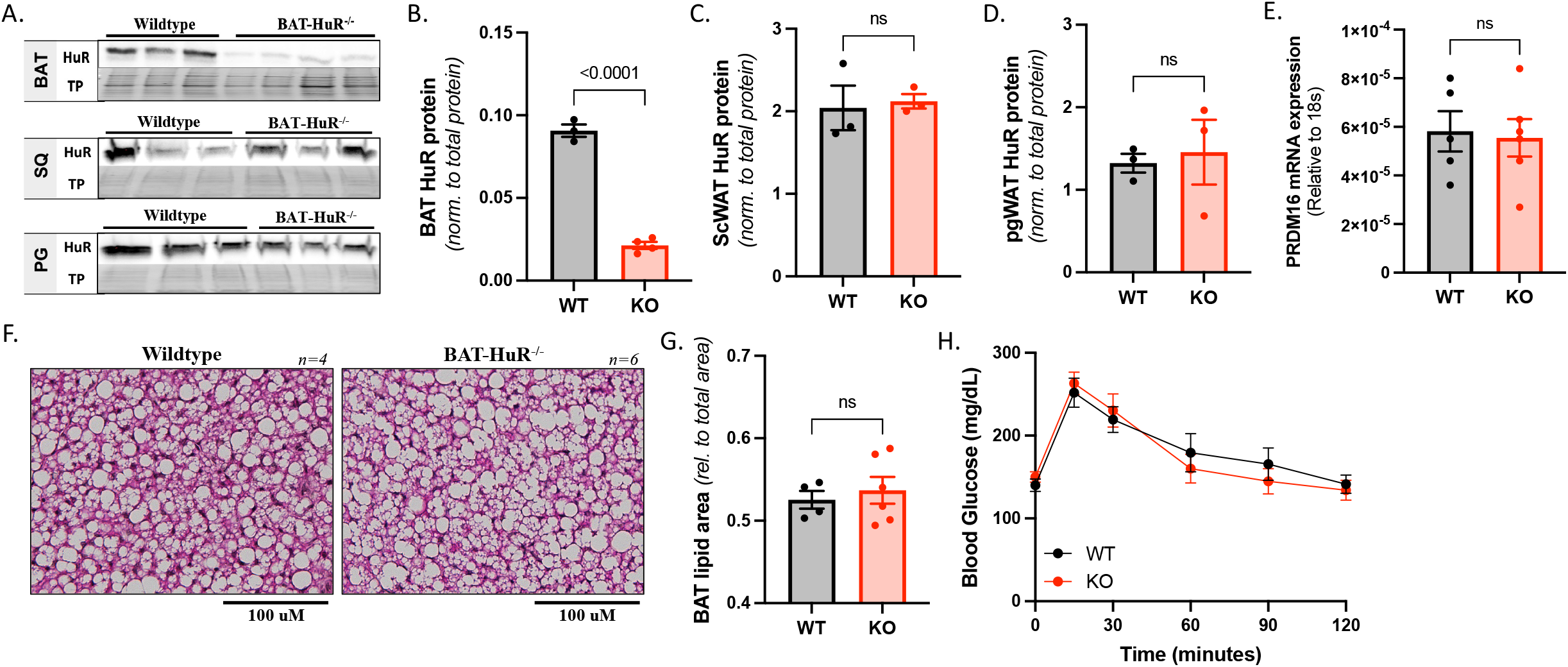
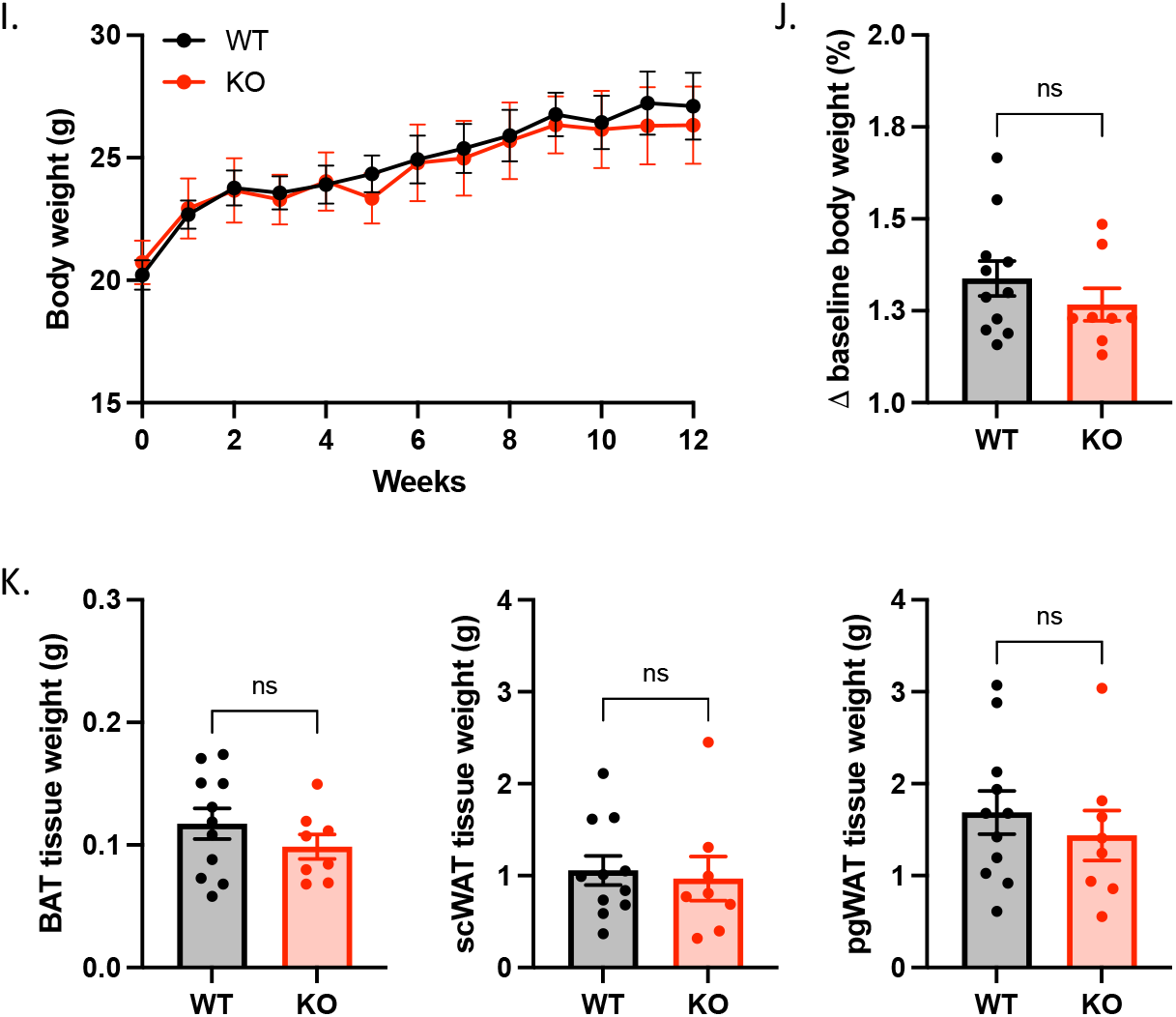
Generation and phenotypic characterization of BAT-specific HuR deletion (BAT-HuR^-/-^) mice. BAT-specific deletion of HuR was achieved by crossing HuR-floxed mice with mice harboring a UCP1-specific Cre recombinase transgene (UCP1-Cre). Western blot of HuR expression in brown (BAT), subcutaneous white (scWAT; SQ), and perigonadal white (pgWAT; PG) adipose tissue show specificity of HuR knockdown to BAT (A; protein expression is quantified in panels B-D). Phenotypically, no differences were observed in BAT expression of PRDM16 mRNA (E), morphology via H&E staining (F), or total lipid droplet area (G). Additionally, BAT-HuR^-/-^ mice showed no difference in glucose tolerance (H), baseline body weight (I; starting weight) or total weight gain following 12 weeks of 45% kcal/fat diet (I-J). We also observed no differences in basal adipose tissue depot weights (K). Data in H is an average of n = 7 wild-type and n = 9 BAT-HuR^-/-^ mice. ns indicates not significant.

Additionally, the baseline body weights of BAT-HuR^-/-^ mice were comparable to wildtype mice and a 12-week high fat diet challenge failed to produce significant differences in overall weight gain (Fig. 1I-J) or the weights of individual adipose tissue depots (Fig. 1K). Together, these results suggest that BAT-HuR^-/-^ mice show no overt adipo-centric phenotype at baseline, suggesting our previously observed lean phenotype of Adipo-HuR^-/-^ mice^23^ is driven by HuR-dependent changes in the WAT.

### BAT-specific deletion of HuR induces an inability to thermoregulate in response to acute cold

To assess the capacity of BAT-HuR^-/-^ mice to thermoregulate under conditions of acute cold, mice were subjected to a cold challenge at 4°C with hourly monitoring of core body temperature (CBT). Relative to WT mice, BAT-HuR^-/-^ mice exhibited a more rapid loss in CBT (Fig. 2A-B) and were also less effective in recovering CBT after a 1h period of return to RT (Fig. 2C). Consistent with our findings in Adipo-HuR^-/-^ mice, we show this effect to be independent of basal or cold-induced UCP1 protein expression (Fig. 2D) or mitochondrial density (Fig. 2E) between WT and BAT-HuR^-/-^ mice.

**Figure 2.**
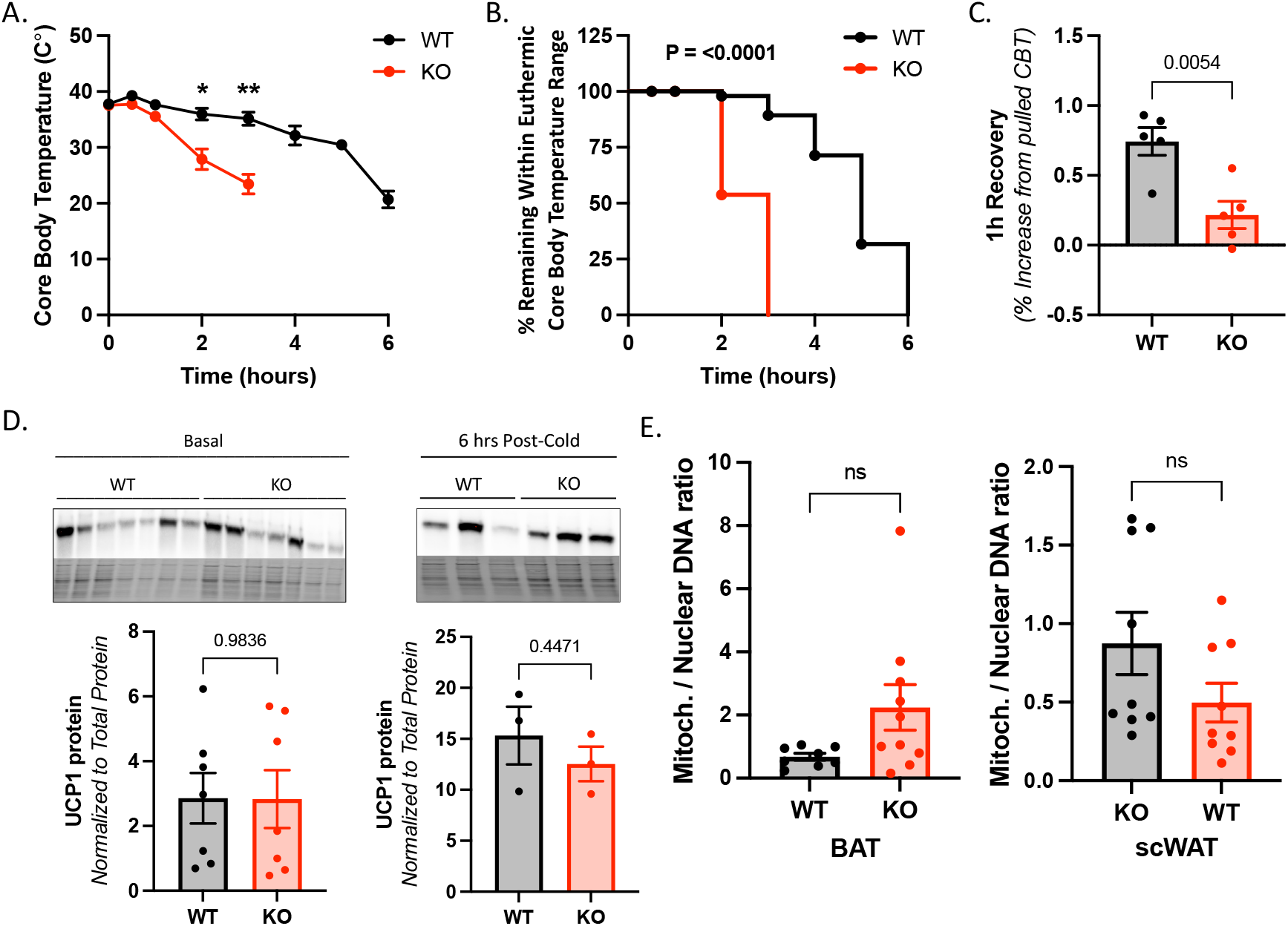
BAT-specific HuR deletion impairs whole body thermogenic capacity in response to acute cold challenge. Core body temperature (CBT) declines more rapidly in BAT-HuR^-/-^ mice upon acute cold challenge at 4°C (A) and BAT-HuR^-/-^ mice were less able to maintain CBT within euthermic range (32-37°C) at 4°C (B). BAT-HuR^-/-^ mice were also slower to recover CBT following removal from 4°C cold challenge (C). Despite being cold intolerant, no difference in UCP1 protein expression levels were observed in BAT-HuR^-/-^ mice at either baseline or at 6 hours post cold challenge (D). The ratio of adipose tissue mitochondrial/nuclear DNA ratio was also unchanged between BAT-HuR^-/-^ and wild-type mice (E). * P ≤ 0.05. ** P ≤ 0.01. ns indicates not significant.

Our previous work revealed BAT-specific decreases in the expression of several calcium transport genes in Adipo-HuR^-/-^ mice.^23^ In addition, sequence analysis of the mRNA transcripts of calcium transport mediators RyR1, RyR2, and SERCA2 identified several putative HuR binding sites within the 3’-untranslated region (3’UTR) of RyR2 and SERCA2 but not RyR1. Moreover, qPCR results confirmed a significant decrease in BAT mRNA expression of RyR2 in BAT-HuR^-/-^ mice (Fig. 3A). However, we did not observe HuR-dependent differences in the mRNA expression of RyR1, SERCA1, SERCA2, or CamKII (Fig. 3B-E). These results further support a BAT-specific role for HuR in regulating calcium-driven thermogenesis, specifically via RyR2.

**Figure 3.**
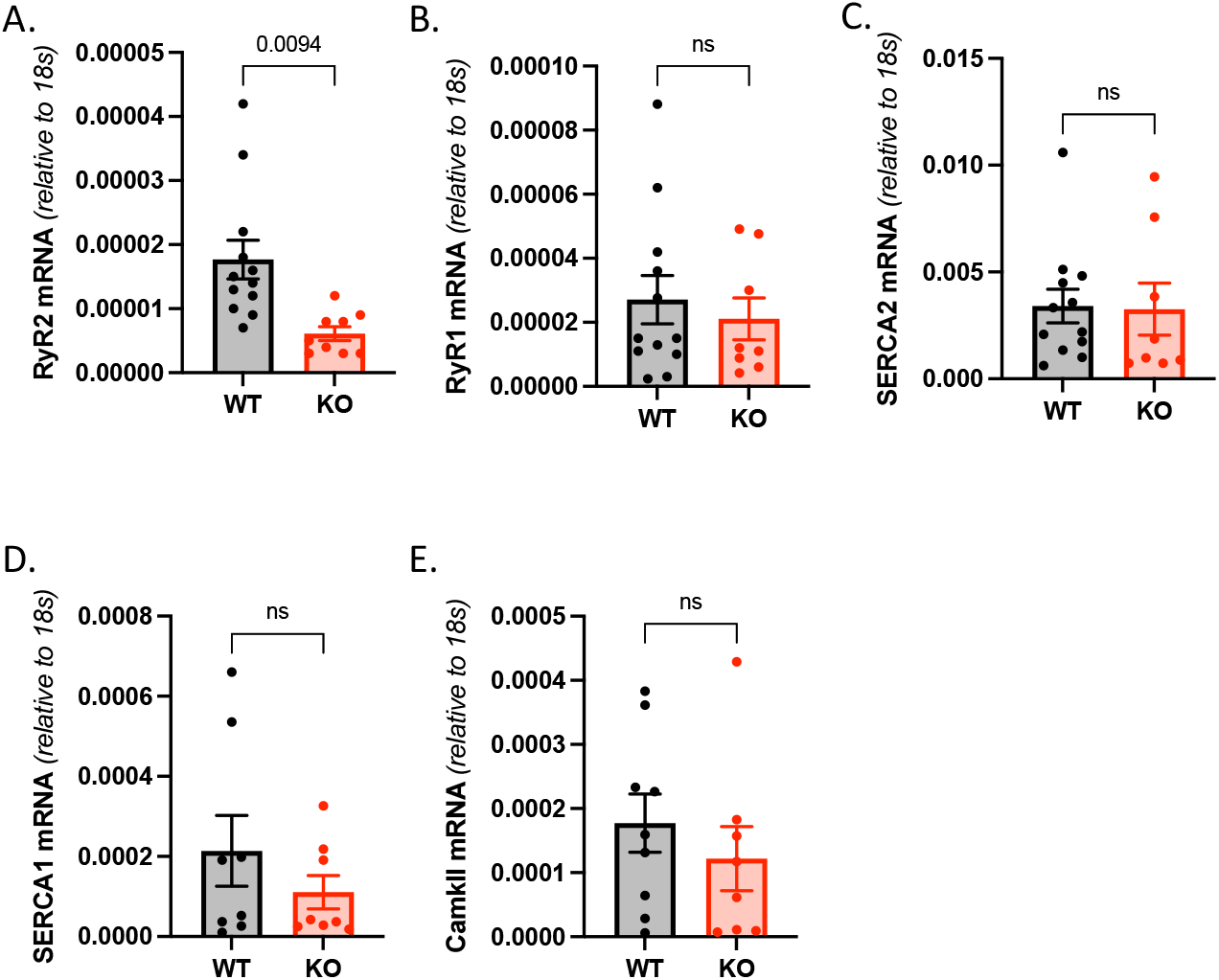
HuR mediates the expression of RyR2, but not additional SR calcium transport-related genes. mRNA expression of SR calcium release genes in BAT from BAT-HuR^-/-^ mice via qPCR shows a significant reduction in RyR2 mRNA compared to wild-type (A), but not RyR1 (B), SERCA2 (C), SERCA1 (D), or CamKII (E).

### Adrenergic-induced cytosolic calcium transport and cellular heat production is HuR-dependent

Since thermogenic metabolism in BAT is driven by β3-adrenergic signaling (β3-AR), we next wanted to assess the direct effect of the β3-AR-specific agonist CL316,243 on HuR activity. Results demonstrate that SVF-derived brown adipocytes (Fig. S1) treated with 10 μM CL show an increase in nuclear-to-cytoplasmic translocation of HuR (Fig. S2), indicative of an increase in HuR post-transcriptional regulatory activity.

To determine the HuR-dependent impact on SR calcium cycling, wild-type (WT) and HuR^-/-^ (KO) SVF-derived primary brown adipocytes were loaded with Fluo-4 AM to assess free cytosolic calcium content and stimulated with the β-AR agonist Isoproterenol (ISO; 10 uM). Our results demonstrate an ISO-induced increase in free cytosolic calcium levels in WT brown adipocytes that is significantly blunted in HuR^-/-^ adipocytes (Fig. 4A-B). A HuR-dependent increase in ISO-mediated cytosolic calcium was recapitulated by similar observations showing a blunting of increased calcium following treatment with DHTS, a pharmacological inhibitor of HuR (Fig. 4C-D). Importantly, this result was found to be maintained upon the depletion of extracellular calcium, demonstrating a specificity for intracellular calcium transport (Fig. S3).

**Figure 4.**
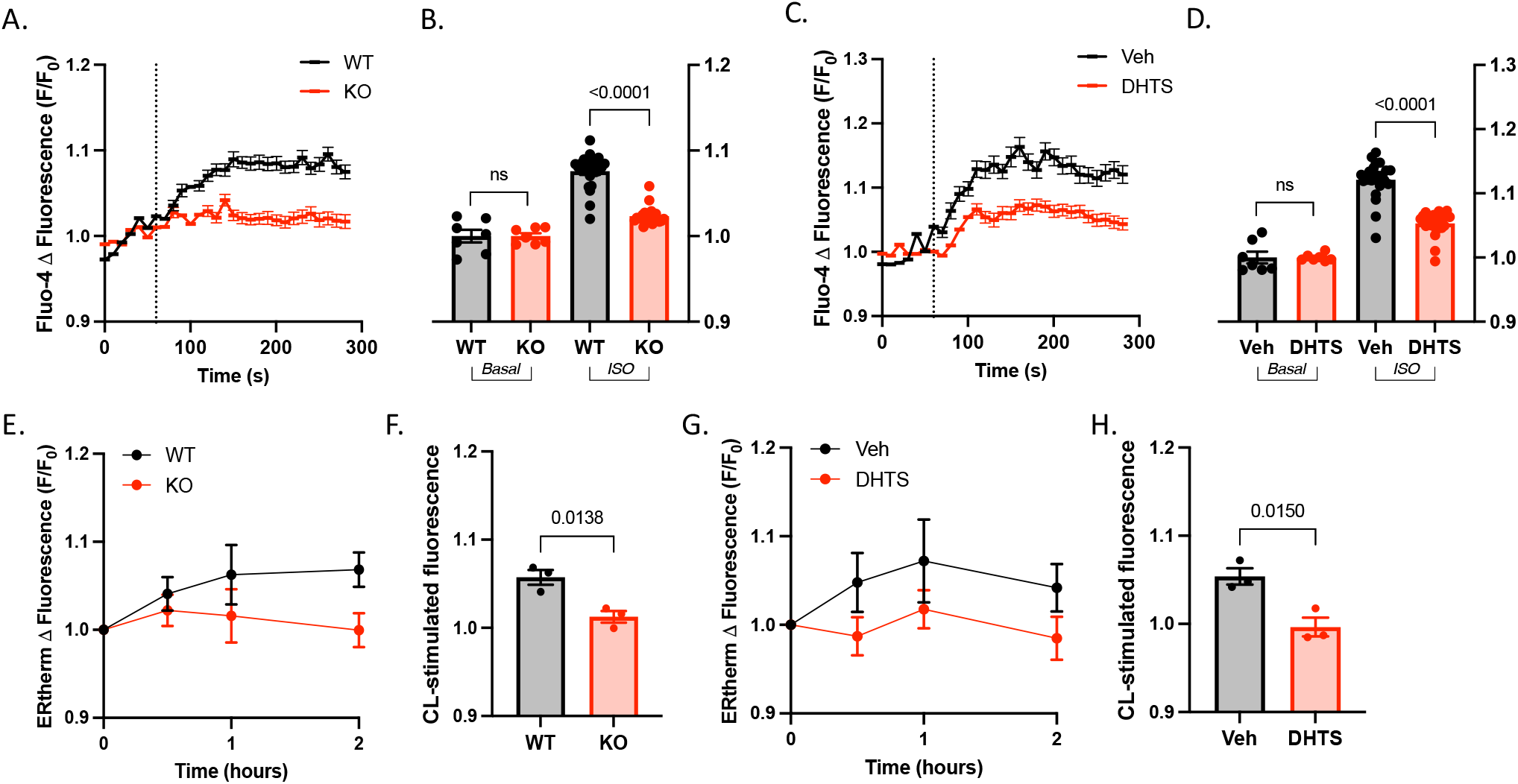
HuR mediates β-AR-dependent increases in cytosolic calcium and thermogenesis in SVF-derived brown adipocytes. β-AR stimulation via isoproterenol (ISO; given at the time indicated by the vertical dashed line) induced an increase in total cytosolic calcium, as determined via Fluo-4, in wild-type cells that is blunted in both HuR^-/-^ cells (A-B) or wild-type cells treated with the HuR inhibitor DHTS (C-D). Calcium response is quantified in panels B and D as total area under the curve (AUC) for individual biological replicates. ERthermAC heat assay revealed a decrease in β3-AR-mediated cellular heat production upon stimulation with 10uM CL316,243 in HuR^-/-^ cells (E-F) and wild-type cells treated with DHTS (G-H). Thermogenic response is measured by a quenching of fluorescence and is represented as the inverse of CL-stiumlated values over basal fluorescent values ([1/(F/F_0_)], and is quantified in panels F and H as total area under the curve (AUC) for individual biological replicates.

We next assessed direct β3-AR stimulated heat production in SVF-derived brown adipocytes using ER-Therm AC fluorescent dye. Results demonstrate an increase in cellular heat following treatment with CL316,243 in brown adipocytes that is blunted in both HuR^-/-^ (Fig. 4E-F) and DHTS-treated cells (Fig. 4G-H). Together, these results show that HuR is necessary for both β-adrenergic-mediated increases in cytosolic calcium concentration and thermogenesis in primary brown adipocytes.

### HuR regulates RyR2 expression in BAT via direct binding and stabilization of RyR2 mRNA

To determine direct HuR binding to RyR2 mRNA, we performed RNA-Protein cross-linking and immunoprecipitation (CLIP) of HuR with its directly bound mRNA targets in wild-type brown adipocytes, followed by RNA isolation and qPCR analysis. We found a significant enrichment in HuR bound RyR2 that was reversed upon DHTS treatment (Fig. 5A). We next sought to determine the mechanistic consequence of HuR binding by assessing the HuR-dependent RyR2 mRNA transcript stability. To do this, cells were treated with actinomycin D (2.5 μg/ml) to inhibit *de novo* transcription followed by treatment with either vehicle or HuR inhibitor, DHTS, for four hours. Results shows a more rapid decrease in RyR2 mRNA in DHTS-treated cells compared to vehicle controls suggesting destabilization of RyR2 mRNA upon HuR inhibition (Fig 5B).

**Figure 5.**
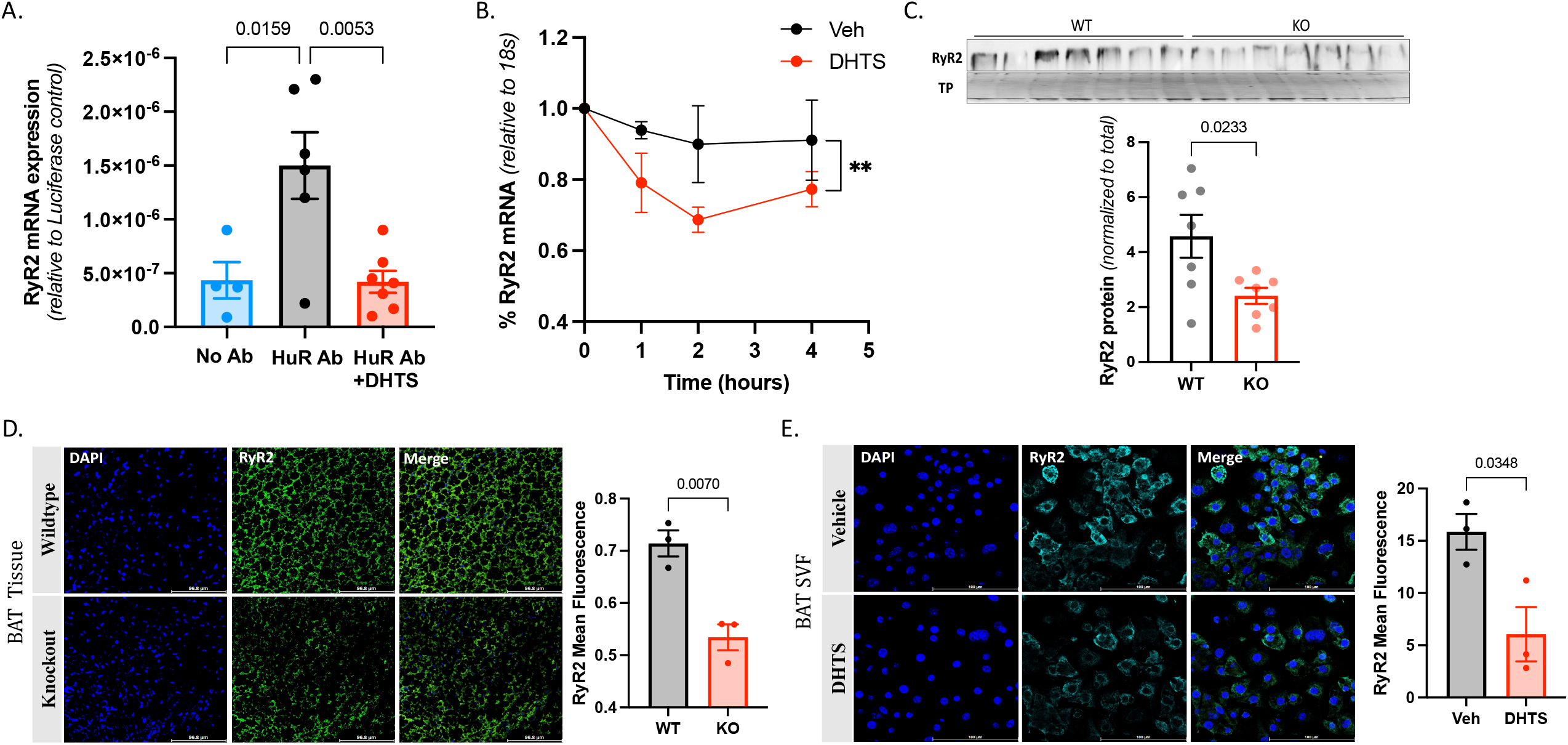
HuR directly binds and stabilizes RyR2 mRNA in brown adipocytes. RNA-protein cross-linking immunoprecipitation (CLIP) of wild-type SVF-derived brown adipocytes reveals a direct HuR binding to RyR2 mRNA, as evidenced by CLIP pulldown with HuR-specific antibody followed by RNA isolation and qPCR detection of RyR2 mRNA, that is mitigated by treatment with DHTS (A). Wild-type SVF-derived brown adipocytes treated with actinomycinD to inhibit transcription and DHTS (or vehicle control) show a more rapid degradation of RyR2 mRNA upon DHTS treatment (B). ** P ≤ 0.01.

Western blotting confirmed a decrease in RyR2 protein in BAT from BAT-HuR^-/-^ mice (Fig. 5C), that was supported by immunofluorescent imaging (Fig. 5D) and DHTS-treated brown adipocytes (Fig. 5E). These results demonstrate that HuR directly binds and regulates the stability of RyR2 mRNA in brown adipocytes and confirms a decrease in RyR2 protein levels upon the loss of HuR expression or activity.

### Rescue of RyR2 activity restores adrenergic-induced cytosolic calcium transport and cellular heat production in HuR^-/-^ brown adipocytes

Wild-type and HuR^-/-^ brown adipocytes were treated with the RyR stabilizer S107 to determine whether enhancing RyR2 activity is sufficient to recover the loss of calcium and heat response observed upon HuR deletion. While S107 did not significantly impact cytosolic calcium in WT adipocytes, it significantly increased the isoproterenol-induced cytoplasmic calcium content of HuR^-/-^ adipocytes (Fig. 6A-B). Similarly, results show a significant increase and full restoration of cellular heat generation in HuR^-/-^ cells, but no impact on WT cells (Fig. 6C-D). These results suggest that deficiency in RyR2 expression and activity is the primary downstream mediator of HuR-dependent calcium-driven thermogenesis in brown adipocytes.

**Figure 6.**
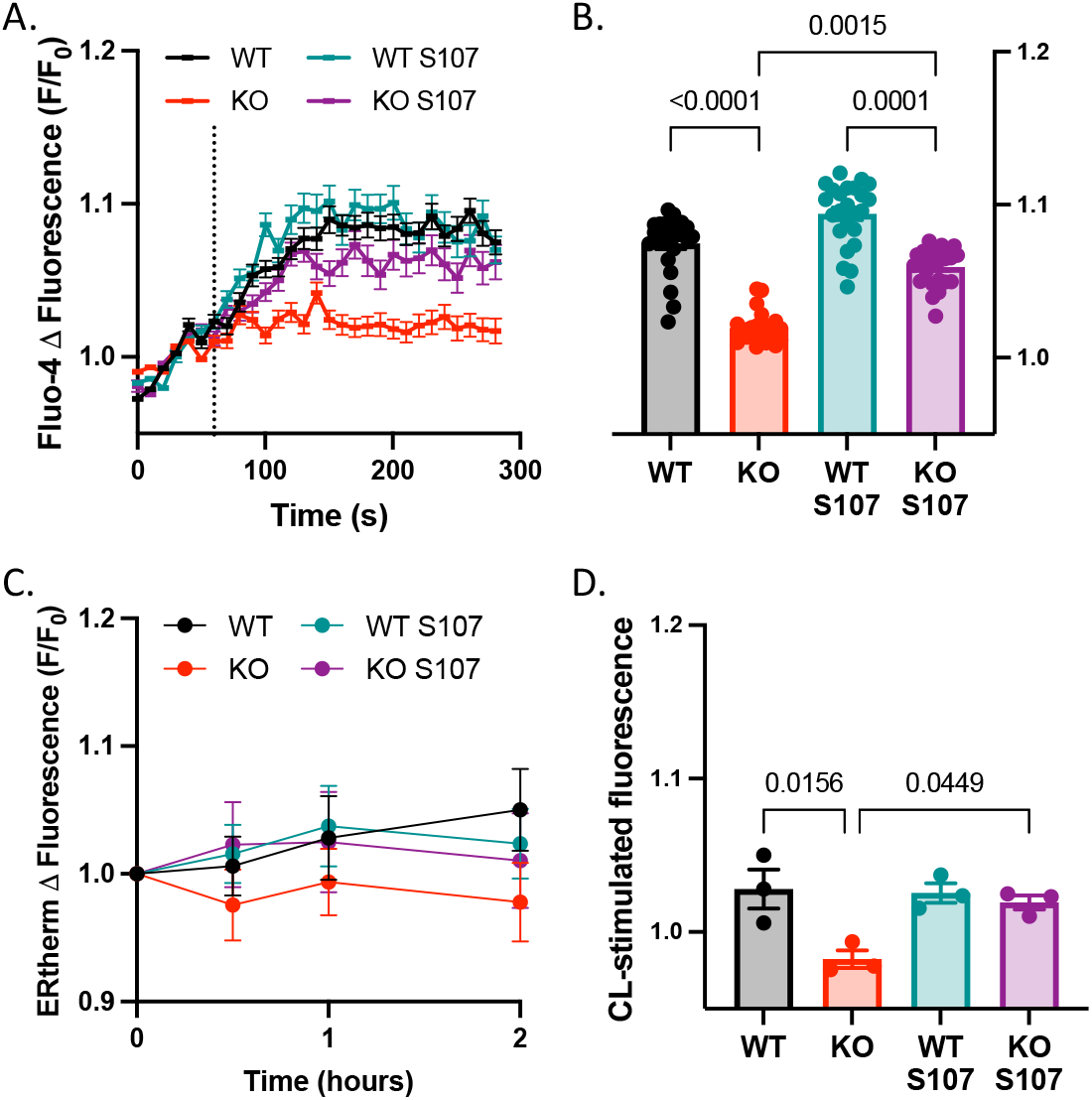
Rescue of RyR2 activity restores adrenergic-induced cytosolic calcium transport and cellular heat production in HuR^-/-^ brown adipocytes. Fluo-4 assessment of total cytosolic calcium shows that S107 restores β-AR-stimulated cytosolic calcium levels (A-B) and cellular heat generation (C-D) in HuR^-/-^, but not wild-type, SVF-derived brown adipocytes.

## Discussion

Obesity is a heterogenous metabolic disease characterized by the accumulation of excess body fat. Despite the recent entry to market of GLP1 receptor agonists for treatment of obesity, these drugs are not appropriate or effective for all patients, and therapies remain limited even in the face of the ongoing rise in obesity rates. Stimulating brown and beige thermogenic adipose tissue activity to upregulate energy expenditure has recently emerged as a therapeutic possibility with the identification of inducible brown adipose tissue in adult humans. Our previous report showed that an adipocyte-specific deletion of RNA binding protein HuR (Adipo-HuR^-/-^) results in an impairment of acute adaptive thermogenesis and suggests that HuR plays a novel role in the thermogenic and functional maintenance of brown adipose tissue through a UCP1-independent pathway.^23^ RNA sequencing results revealed a decrease in several SR calcium transport genes, suggesting reduced SR calcium cycling as a potential mechanism for the observed thermogenic disruptions.^23^ Here, we experimentally show a direct role for HuR in mediating SR calcium cycling and calcium-dependent thermogenesis through regulation of RyR2 expression in brown adipocytes.

The results presented herein describe a post-transcriptional regulatory relationship between HuR and SR calcium mediator RyR2 that is required for BAT participation in calcium-driven thermogenesis. Since HuR has previously been suggested to regulate adipocyte differentiation, we generated mice with a BAT-specific deletion of HuR (*BAT-HuR*^*-/-*^) driven by a UCP1-Cre promoter to induce genetic deletion of HuR in mature brown adipocytes. Interestingly, while we reported reductions in body weight and BAT lipid content at baseline in chow-fed Adipo-HuR^-/-^ mice, BAT-HuR^-/-^ showed no difference in baseline body weight or lipid content as compared to controls. However, the cold sensitive phenotype reported in *Adipo-HuR*^*-/-*^ mice was maintained in *BAT-HuR*^*-/-*^ mice acutely challenged by cold, confirming that HuR expression is critical to maintaining thermogenic adipocyte function and loss of HuR expression in BAT is sufficient to induce whole-body thermal impairment. Consistent with our previous findings, BAT-HuR^-/-^ mice did not show any differences in UCP1 protein expression or mitochondrial density in BAT or scWAT depots indicating the cold sensitive phenotype is likely independent of canonical UCP1 thermogenic signaling and mitochondrial ATP production. Thus, BAT-HuR^-/-^ mice are a more suitable model to determine the mechanistic underpinnings of HuR-dependent thermogenesis in BAT.

Since HuR deletion is driven by cre-recombinase under control of the UCP1 promoter, recombination should only be observed in mature brown adipocytes, reducing the chance of compensatory effects due to the loss of HuR expression. In addition, the mRNA expression of brown adipocyte differentiation mediator PRDM16 was unchanged, suggesting adipogenesis and brown adipose tissue differentiation programs were unaffected in this model. Moreover, replication of our results using pharmacological inhibition via DHTS provides further evidence for the acute HuR specificity of our observations. Additionally, it should be noted that since UCP1 is also expressed in beige adipocytes within select WAT depots, we can’t conclusively rule out a HuR-dependent contribution in beige adipocytes to our *in vivo* cold challenge data shown in Fig 2. Nonetheless, our *in vitro* studies show a clear role for HuR in modulation of calcium-dependent thermogenesis in brown adipocytes.

Stimulation of α1 and β-adrenergic receptors with norepinephrine was previously demonstrated to increase intracellular calcium signaling in beige adipocytes through SR-calcium release, thus subsequently enhancing SERCA2b ATP consumption as it pumps calcium back to the SR.^24^ The stimulation of mitochondrial ATP production by elevated cytoplasmic calcium, along with the concordant consumption of ATP by SERCA, is responsible for energy dissipation in the form of heat.^25^ Herein, we report the activation of HuR in BAT, evidenced by its nuclear to cytoplasmic translocation, downstream of β3-adrenergic agonism via CL316,243. We further show that the increase in cytosolic calcium, presumed to be due to SR calcium release, in response to β-adrenergic stimulation is decreased in HuR^-/-^ and DHTS-treated brown adipocytes. The primary sources of cytosolic calcium are likely to be extracellular calcium influx by L-type calcium channels (LTCC) or SR calcium release by RyR or inositol 1,4,5-trisphosphate (IP3) receptors. However, deficiencies in cytosolic calcium were maintained when cells were placed in calcium-free buffer indicating the involvement of HuR in coordinating intracellular calcium release from the SR.

We hypothesized that HuR regulates the mRNA stability of RyR2 based on our previously published RNA-sequencing data on BAT from Adipo-HuR^-/-^ mice and the presence of multiple canonical AU-rich HuR binding sites within the RyR2 3’UTR. In addition, previous work has shown direct binding and regulation of RyR2 activity by highly conserved Hu-family member HuD (*elavl4*) in motor neurons.^26^ HuR is a ubiquitously expressed member of Hu-family proteins, all of which contain three highly conserved RNA recognition motifs sharing >90% amino acid sequence identity. The prominent function of the Hu-family members is direct binding to the 3′ UTR of target mRNAs, most typically resulting in enhanced mRNA stability and translation to promote an increase in protein expression. Upon observing decreased mRNA and protein expression of RyR2 in BAT-HuR^-/-^ mice, we further demonstrate direct HuR binding to and stabilization of RyR2 mRNA in brown adipocytes.

Our results further demonstrate a key role for HuR-dependent expression of RyR2 and define the functional contribution of RyR2 in HuR-dependent calcium release through a ‘rescue’ of SR calcium release specifically in HuR deficient cells using the RyR-Calstabin interaction stabilizer, S107. S107 treatment partially restored β-adrenergic-mediated cytosolic calcium levels of BAT-HuR^-/-^ adipocytes, but did not further enhance SR calcium release in wild-type cells. The inability of S107 to further enhance the cytosolic calcium response in WT cells could be attributed to an already near-maximal activation of RyR calcium release activity upon isoproterenol stimulation in functional cells. Further, S107 has been shown to mitigate pathological calcium leakage in myocytes, and therefore could also function to potentiate SR calcium loading and maintain the balance between cytoplasmic and SR-sequestered calcium.^27–29^ Functionally, S107 treatment was also able to almost fully restore β-adrenergic-mediated thermogenesis in HuR^-/-^ cells to that of wild-type levels. This result is interesting and suggests a pool of spare RyR2 channels and that less than 100% of the basal RyR2 pool is sufficient to mediate maximal calcium driven thermogenesis in brown adipocytes. This data also fundamentally demonstrates that brown adipocytes do in fact participate in calcium-driven thermogenesis as an increase in cytosolic calcium resulted in increased heat generation in BAT-HuR^-/-^ brown adipocytes.

Importantly, we show that the HuR-dependent loss of thermal regulation at the whole organism level is translatable in mechanistic studies in primary brown adipocytes, and suggests that the homeostatic maintenance of SR calcium cycling impacts thermal output at the cellular level. It is important to note that they possess a lower capacity for ATP synthesis compared to beige adipocytes who generate ATP through glycolysis and participation in TCA and electron transport cycle processes.^30^ It has been suggested that calcium-driven thermogenesis by action of SERCA ATP hydrolysis is dispensable in BAT, but not beige adipose tissue, in the presence of UCP1.^24^ However, this does not necessarily mean that this thermogenic pathway is unique to beige fat, and we now clearly show a calcium-dependent thermal output at the cellular level in brown adipocytes. Taken together, these results more likely indicate a greater reliance of beige adipose tissue on non-canonical thermogenic mechanisms due to its more limited UCP1 expression compared to BAT. A noted limitation of this work is the inability to conclusively determine the tissue-specific contribution of brown adipose involvement in calcium-driven thermogenesis on whole-body thermal metabolism. Thus, it is possible the defects in thermal capacity observed in BAT-HuR^-/-^ mice are driven in part by impaired calcium homeostasis in beige adipose tissue. While a potential role for HuR in calcium-mediated thermogenesis in beige adipocytes would need to be delineated by future studies, our results clearly show a functional role for HuR-mediated RyR2 expression in brown adipocytes through quantifiable differences in thermal activity at the cellular level.

## Acknowledgements

Fluorescent microscopy was done with the support of Chet Closson at the University of Cincinnati Live Microscopy Core. The Leica Stellaris support from the Office of the Director, National Institutes of Health under Award Number S10OD030402. The content is solely the responsibility of the authors and does not necessarily represent the official views of the National Institutes of Health.

