## Supplementary figures and images for "HuR-dependent expression of RyR2 contributes to calcium-mediated thermogenesis in brown adipocytes"

### Supplemental Figures

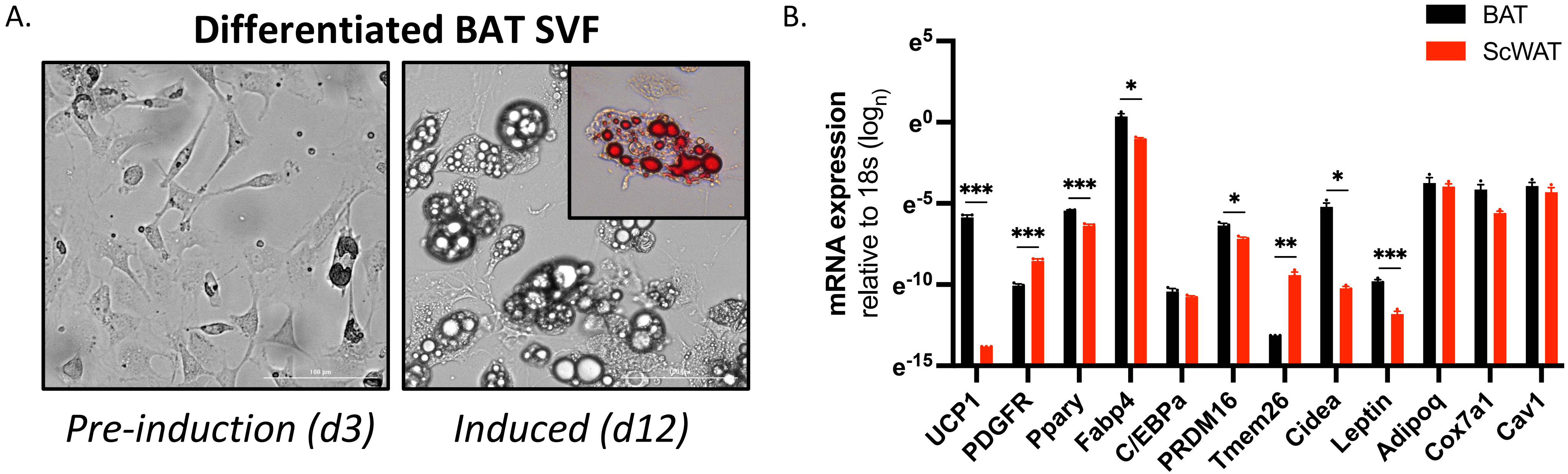

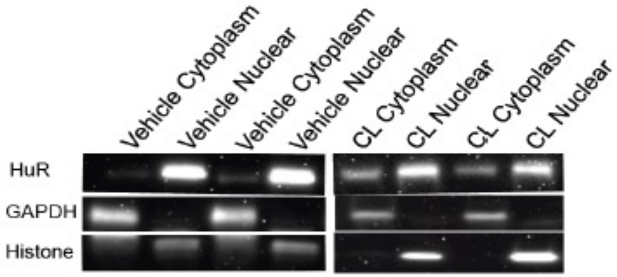

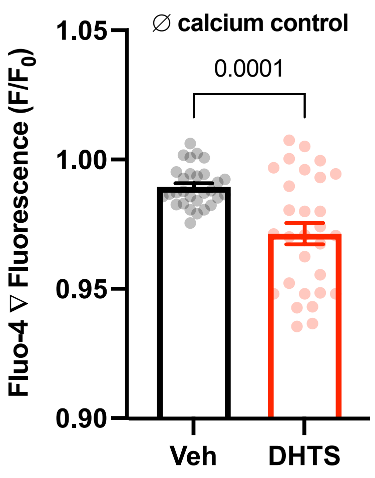
